# Risk of mitochondrial deletions is affected by the global secondary structure of the human mitochondrial genome

**DOI:** 10.1101/603282

**Authors:** Victor Shamanskiy, Alina A. Mikhailova, Kristina Ushakova, Alina G. Mikhailova, Sergei Oreshkov, Dmitry Knorre, Evgenii O. Tretiakov, Natalia Ri, Jonathan B. Overdevest, Samuel W. Lukowski, Irina Gostimskaya, Valerian Yurov, Chia-Wei Liou, Tsu-Kung Lin, Wolfram S. Kunz, Alexandre Reymond, Ilya Mazunin, Georgii A. Bazykin, Konstantin Gunbin, Jacques Fellay, Masashi Tanaka, Konstantin Khrapko, Konstantin Popadin

## Abstract

Aging in postmitotic tissues is associated with clonal expansion of somatic mitochondrial deletions, the origin of which is not well understood. Deletions in mitochondrial DNA (mtDNA) are often flanked by direct nucleotide repeats, but this alone does not fully explain their distribution. Here, we hypothesized that the close proximity of direct repeats on single-stranded DNA might play a role in the formation of deletions. By analyzing human mtDNA deletions in the major arc of mtDNA, which is single-stranded during replication and is characterized by a high number of deletions, we found a non-uniform distribution with a "hot spot" where one deletion breakpoint occurred within the region of 6-9kb and another within 13-16kb of the mtDNA. This distribution was not explained by the presence of direct repeats, suggesting that other factors, such as the spatial proximity of these two regions can be the cause. In silico analyses revealed that the single-stranded major arc may be organized as a large-scale hairpin-like loop with a center close to 11kb and contacting regions between 6-9 kb and 13-16 kb, which would explain the high deletion activity in this contact zone. The direct repeats located within the contact zone, such as the well-known common repeat with a first arm at 8470-8482 bp and a second arm at 13447-13459 bp, are three times more likely to cause deletions compared to direct repeats located outside of the contact zone. An analysis of age- and disease-associated deletions demonstrated that the contact zone plays a crucial role in explaining the age-associated deletions, emphasizing its importance in the rate of healthy aging. Overall, we provide topological insights into the mechanism of age-associated deletion formation in human mtDNA, which could be used to predict somatic deletion burden and maximum lifespan in different human haplogroups and mammalian species.

## Introduction

Aging is associated with the accumulation of DNA damage. The mitochondrial genome (mtDNA), existing within a cell in a large number of copies, is strongly predisposed to accumulate such age-related damage due to continuous turnover [1] and high mutation rate [2]. The coexistence of different mtDNA variants within the same cell (heteroplasmy) [3] assures an intracellular mtDNA competition, which is especially influential in slowly-dividing tissues, where the “selfish” mtDNA mutants with replication advantage but functional disadvantages have a time for clonal expansion [4]. One of the best-studied examples of selfish mtDNA mutations is deletions - the elimination of a portion of a mitochondrial genome. In *substantia nigra* neurons, for example, the first mtDNA deletions were detected at around 50 years of age [4,5]. Each year, this fraction of heteroplasmy increased by 1-2% until, after several decades, a phenotypically essential threshold of 50-80% was reached [6], leading to neurodegeneration. Skeletal muscle is another tissue that is predisposed to the accumulation of mtDNA somatic deletions: an expansion of somatic mtDNA deletions within myofibrils is associated with sarcopenia - loss of muscle weight and strength with age [7,8]. Other tissues with slow-dividing cells that are also affected by mtDNA deletions include extraocular muscles [9] and oocytes [10–12]. In the case of oocytes, the expansion of mtDNA deletions could potentially manifest itself across all tissues, including proliferative ones, leading to multisystem disorders [13,14].

There are several lines of evidence supporting the hypothesis of somatic mtDNA deletions causing host cell degeneration and several corresponding age-related phenotypes. (i) The proof-reading-deficient version of mtDNA polymerase leads to the accumulation of somatic point mutations and deletions in mice, followed by reduced lifespan and premature onset of aging-specific phenotypes ([15][16]). However, the level of point somatic mutations is rather low in normal mice questioning the role of mtDNA mutations in normal aging [17,18]. (ii) The observation of localization of mtDNA deletions in regions of myofiber breakage [19] and respiratory chain-deficient neurons [4] supports the hypothesis of the causative effects of mtDNA deletions on aging. (iii) A reported deficit of neurons carrying an extremely high (> 80%) deletion burden suggests that such cells are degraded and no longer present in the analyzed tissue [6]. Altogether, the high mtDNA deletion burden is not a neutral hallmark of aged cells but is more likely a causative agent. Thus, understanding the molecular mechanisms underlying the origin of somatic mtDNA deletions, as well as their rate of expansion, is of primary importance [20,21].

It has been shown that most somatic mtDNA deletions are flanked by direct nucleotide repeats [22] or by long imperfect duplexes consisting of short stretches of direct repeats [23]. Since direct repeats predispose mtDNA to somatic deletions, they are considered to be an example of “Deleterious In Late Life” alleles (DILL): neutral or slightly-deleterious during reproductive age but harmful in late life [24]. The negative correlation between the abundance of direct repeats in mtDNA and the species-specific lifespan of mammals [25,26] has been interpreted as additional evidence of the deleterious effect of repeats in mtDNA of long-lived mammals. Similarly to a deficit of direct repeats in mtDNA of long-lived mammals, we previously hypothesized that the decreased number of direct repeats in the mitochondrial genome of some human haplogroups could be linked to a lower prevalence of somatic mtDNA deletions, thereby enabling healthier aging and postponing the aging process [27].

Although the direct nucleotide repeats (or long imperfect duplexes) have long been known to contribute to the formation of mtDNA deletions, they still only explain a small fraction of observed deletion distributions. This raises questions about why some repeats lead to deletions while others do not, and what other factors may be involved in deletion formation. To understand the main factors behind mtDNA deletion formation, it is reasonable to start with an analysis of the most mutagenic direct repeat in the human genome: the common repeat ([22], [23], [27]) which is the longest perfect direct repeat in the human mtDNA. An important feature of the common repeat is that its “arms” (first arm at 8470-8482 bp and a second arm at 13447-13459 bp) are located exactly in the peaks of the distribution of all deletion breakpoints across mtDNA. Based on this observation, it has been hypothesized that the common repeat “appears to be the principal factor behind the formation of most deletions” [22]. It means that the common repeat may be important not only for the formation of the common deletion but also to play a role in the emergence of all other mtDNA deletions. To test this hypothesis, we analyzed the mtDNA deletion spectrum in the frontal cortex of samples from the N1b1 haplogroup where the repeat was disrupted (the proximal arm was acTtccctcacca versus acCtccctcacca as it is in the vast majority of the human population). If there was a special structural role of the common repeat, we expected to see that the distribution of all mtDNA deletions within N1b1 samples would differ from other haplogroups with the perfect common repeat. Within our sample size (two cases and two controls), we observed a near complete absence of the common deletion *per se* in N1b1 samples, however, we didn’t find any changes in the distribution of other deletions [23]. Thus we rejected the hypothesis that the common repeat is the main factor behind the formation of the majority of deletions [22]. Rejection of this hypothesis left the main observation, emphasized by Samuels and coauthors, unexplained - i.e. why is the distribution of deletions within the major arc strongly non-uniform? This non-uniformity in the distribution of deletions requires a novel explanation.

Here, by drawing parallels between deletions in bacteria [28], mtDNA common repeat [29], and nuclear DNA [30], we hypothesized that direct repeats might be more likely to cause deletions when they are in close proximity to each other. Thus the increased probability of deletions appearing near the common repeat is maintained not by the common repeat per se but by an independent topological factor. In order to test this hypothesis, we reconstructed the potential spatial structure of the single-stranded major arc of mtDNA and proposed that the major arc is organized as a large-scale hairpin-like loop with a center close to 11 kb and a stem between 6-9 kb and 13-16 kb. This infinity-symbol shape of mtDNA affects the mutagenic potential of direct repeats and thus shapes the distribution of deletions.

## Results

### 1. The deletion spectrum is non-uniform and poorly explained by the direct repeats

If the formation of deletions depends on the spatial proximity of single-stranded DNA regions, we expect that the distribution of deletions will be non-uniform and will follow the structure of the DNA. To understand the potential structure of single-stranded DNA regions, we analyzed the distribution of deletions within the major arc of human mtDNA, where most deletions occur. To do this, we used data from the MitoBreak database, which contains a collection of human mtDNA deletions [31]. We examined the distribution of the centers of each deletion within the major arc. We found that the median center was located at 11,463 bp (Fig 2A, right vertical panel, N=1060), and the distribution was relatively narrow, indicating that there is a low variation in the position of centers and they tend to cluster together. To confirm this, we compared the observed variation in the position of centers with randomly generated ones (see methods). Our analysis revealed that the observed variation was indeed significantly lower than expected (p-value < 0.0001). The observation that most deletions are narrowly clustered around 11,463 bp suggests that a single-stranded major arc can be folded into a hairpin-like structure with a folding axis around 11,463 bp (see several potential schemes in Fig 1).

**Figure 1.**
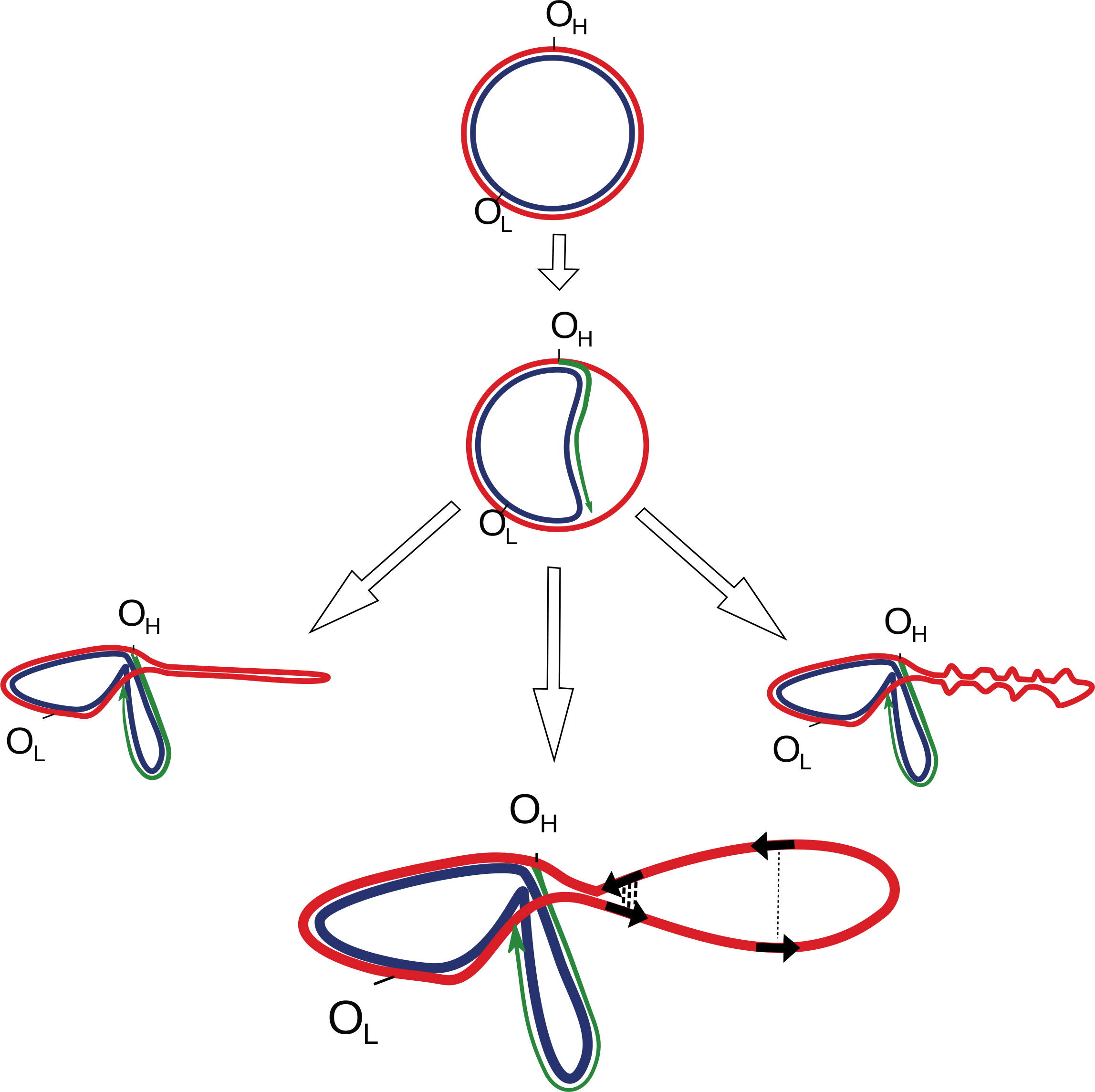
Potential secondary structures formed by a single-stranded parental heavy chain during mtDNA replication. The lower panel shows that direct repeats, marked by black arrows, have different chances of being realized into deletions as a function of a spatial structure. The close spatial proximity of repeats (bold dotted lines) increases the probability of deletion formation, while for repeats that are spatially separated by a greater distance, this probability is decreased (thin dotted line).

**Figure 2.**
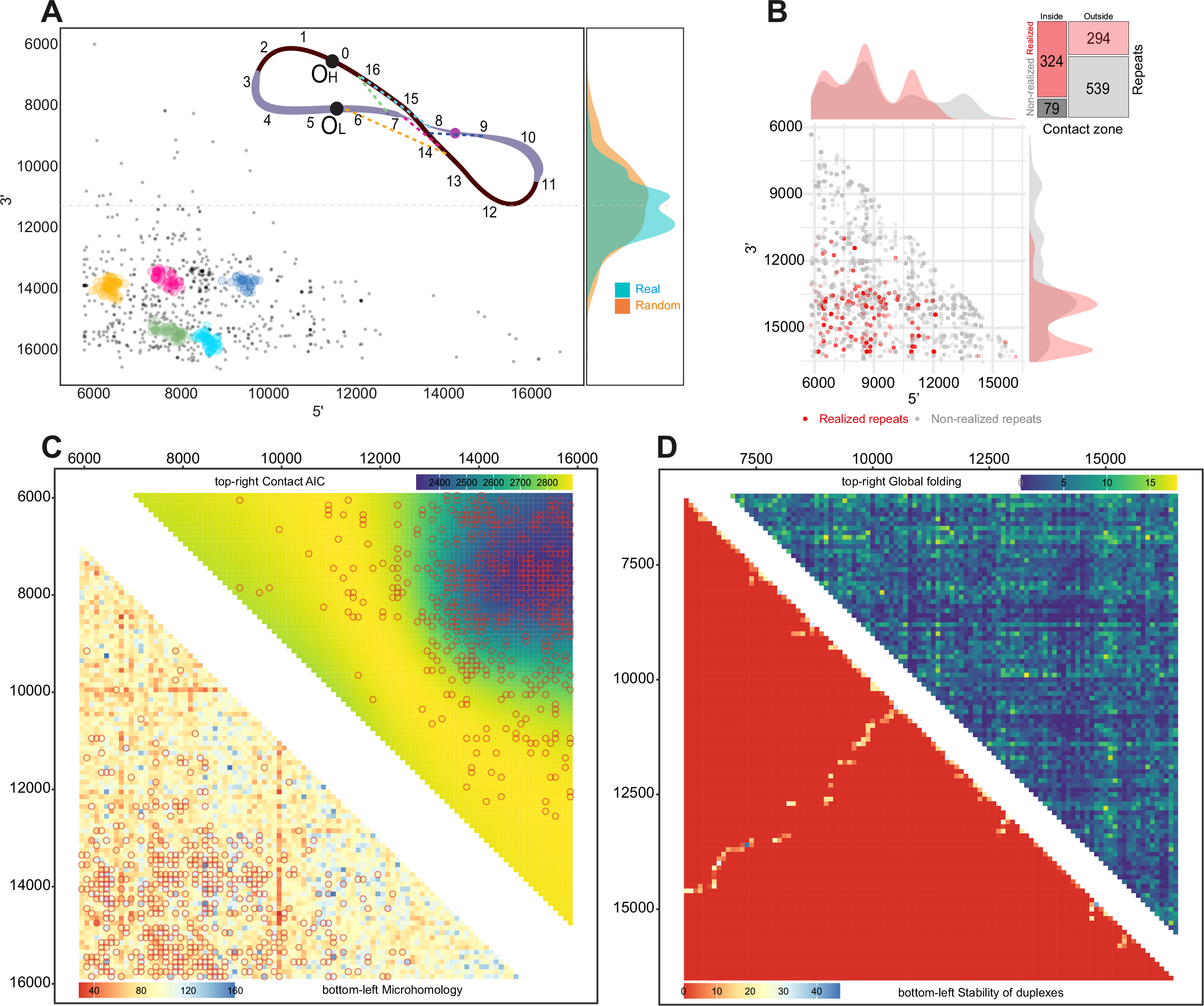
Secondary structure of mtDNA. **A**. Clusters of deletions within the major arc. The majority of clusters are located close to each other within the potential contact zone. The colors on the scheme on top correspond to the clusters. The vertical density plots on the right part of the figure demonstrate the distribution of deletion centers: real (observed) and random (expected); **B.** Realized (red) versus non-realized (gray) repeats tend to be enriched in the potential contact zone. Mosaic-plot of repeats (realized versus non-realized) within and outside the potential contact zone; **C.** Bottom-left: heatmap of the microhomologies between 100 bp windows within the major arc. Microhomology alone poorly explains the distribution of the deletions (empty circles). Top right: heatmap of the data-driven contact zone, based on the AIC of the compared models. **D.** *In silico* approach for major arc global folding prediction. Bottom left triangle: contact matrix derived from the *in silico* folding estimation of the major arc's whole single-stranded heavy chain; top right triangle: contact matrix derived from the *in silico* folding estimation of 100 bp windows of the single-stranded heavy chain of the major arc.

Deletion breakpoints: 3’ and 5’ coordinates are expected to be more abundant in the regions, spatially proximate to each other. To reveal the potential non-uniformity in the distribution of breakpoints, we grouped individual deletions into clusters (Fig 2A clusters are presented by coloured dots, see Methods). We observed that the largest clusters are located close to each other in a specific region, with 5’ breakpoints between 6-9 kb and 3’ breakpoints between 13-16 kb (Fig 2A). This suggests, that a single-stranded major arc forms a stem where 6-9 and 13-16 kb are spatially close to each other; the deficit of breakpoints outside of this region (9-13kb) suggests that this section of the single-stranded major arc can be maintained as an open loop. Importantly, an approximate center of this loop (11 kb) is consistent with the predicted folding axis (11.463 bp) from the analyses of centers (Fig 2A right vertical panel; see also a scheme of the mtDNA at the right-top part of Fig 2A).

During the asynchronous replication of mtDNA, the parental heavy strand of the major arc gradually becomes single-stranded. The region closest to the origin of replication of the heavy strand (with approximate coordinates 16,5 kb) is an early-replication region, which becomes single-stranded first, and as the replication fork moves, the entire major arc becomes single-stranded with the last region close to the origin of replication of the light strand (with approximate coordinates 6 kb). If the time being single-stranded is important for deletion formation, as it has been suggested for the mutagenesis of single-nucleotide substitutions [32,33], we can assume that the early-replicating region (~ 16.5 kb) may be more mutagenic as compared to the late-replicating region (~ 6 kb). The increased deletion mutagenicity of the early-replicating regions (16.5 kb, which corresponds to 3’ breakpoint) as compared to the late-replicating region (6 kb, which corresponds to 5’ breakpoint) can be realized in the fact that the early-replicating region is less selective: this region can associate with any other open regions of the major arc, meaning an increased variation in 5’ deletion breakpoint as compared to 3’ deletion breakpoint. Analysis of a scatterplot of deletion breakpoints and clusters (Fig 2A) confirms this, showing that colored clusters are better described as an oval with increased length along the x-axis, indicating increased variation in 5’ breakpoints. Overall, the increased variation in 5' breakpoints compared to 3' breakpoints (Fig 2A) suggests that deletion formation may also be affected by the amount of time a strand is single-stranded: this is higher for 3' breakpoints, allowing them to associate with a wider range of 3' breakpoints.

All the above-mentioned results suggest that the single-stranded major arc can fold into a large loop with a center close to 11.5 kb and a stem formed by regions 6-9 and 13-16 kb, where the early-replicated 13-16 kb part of a stem can associate with a broad range of late-replicated regions: not only 6-9 kb but also 10 and 11 kb for example. However, till now all our analyses were agnostic - without considering the information that the distribution of deletions is partially explained by direct repeats within the human mtDNA [22,23] and bacterial genomes [28]. To test the importance of the spatial DNA structure as a factor affecting the formation of the deletions, we have to take into account direct repeats also. To do it, we compared the distribution of the perfect direct repeats (see Methods) and deletions from the MitoBreak database. While direct repeats can explain the local distribution of deletions within specific regions, such as the 6-10 versus 12-16 kb region [23], globally, on the scale of the entire major arc, they have a poor correlation with the distribution of deletions. Indeed, we observed an approximately uniform global distribution of the direct repeats within the major arc versus the strongly biased distribution of the deletions (Supplementary Fig 1). This observation is in line with the previous finding by Samuels et al [22] and confirms that direct repeats alone do not fully explain the distribution of deletions in the mtDNA and highlights the need to consider other factors such as the role of spatial DNA structure in the formation of mtDNA deletions.

Deletions may be caused by specific, for example, C-rich motifs [29] within the direct repeats. Thus, there is a possibility that the shifted distribution of deletions might be explained by the shifted distribution of such motifs - for example, hot, deletion-induced motifs can be located preferentially in 6-9 and 13-16 kb regions. In order to test the potential impact of specific motifs within direct repeats on the formation of deletions, we analyzed our database of degraded repeats of the human mtDNA [34], grouping them according to motifs. We then combined all repeats with the same motif into all possible pairs where one repeat was "realized" (if there is a MitoBreak deletion flanked by these nucleotide sequences) and another was "non-realized" (if there is no corresponding deletion in MitoBreak). Our comparison of the positions of realized and non-realized repeats demonstrated that realized repeats were more likely to be located near the 6-9 and 13-16 kb region, while non-realized repeats were more uniformly distributed throughout the major arc (Fig 2B). Motif-specific paired analyses, where we compared properties of realised versus non-realised repeats with the same motif, revealed that non-realized repeats tend to start 700 bp later and end 1300 bp earlier, resulting in a 2000 bp shorter distance between arms of non-realized repeats compared to realized repeats (all p-values < 2.2e-16, paired Mann-Whitney U-test; Number of clusters = 618).

Overall, we observed that the distribution of deletions is highly non-uniform (Fig 2A,) and this non-uniformity is not linked to either direct repeat abundance (Supplementary Fig 1) or direct repeat motifs (Fig 2B). Taking into account that 80% of realized repeats start in the interval 6465-10954 bp and end at the interval 13286-15863 bp, we suggest that this biased distribution can be explained by the potential macromolecular contact zone of the single-stranded DNA between 6-9 and 13-16 kb. Indeed, there is a strong excess of realized repeats within the 6-9 and 13-16 kb region (Fig 2B, mosaic-plot; Fisher odds ratio = 7.5, p < 2.2e-16).

### 2 Probability of deletions is a function of both DNA microhomology and the proximity to the contact point

We have shown that the distribution of the deletions within the major arc is poorly explained by the distribution of direct repeats alone while the potential global structure of the single-stranded mtDNA can be an extra factor affecting deletion formation (Fig 1, Fig 2A, 2B). Here, we aim to build a multiple model including both repeats and secondary structure as major factors affecting deletion formation. Instead of direct repeats, we derived a more biologically meaningful “microhomology similarity” metric, which is (i) an integral metric of similarity between two regions of DNA and (ii) it is fixed to 100 bp windows to facilitate downstream analyses (see Methods). We estimated all pairwise microhomology similarities between all 100 bp windows inside the major arc (Fig 2C bottom - left triangle) and, first of all, as expected, obtained a positive correlation between the microhomology similarity and the density of direct repeats in the corresponding windows (see Methods, Spearman’s Rho = 0.07, P = 1.698e-06, N = 4950 regions 100 bp x 100 bp windows). Next, to analyze an association between deletions and the microhomology similarity, we performed logistic regression where the presence or absence of deletions in each 100×100 bp cell (coded as 1 if there is at least one deletion in a cell, N = 484; coded as 0 if there are no deletions in a cell, N = 4466) was estimated as a function of the Microhomology Similarity (MS):

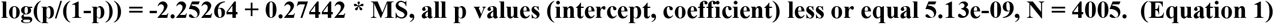

This result confirms our previous findings [23] and shows that a high microhomology similarity is positively correlated with a higher probability of deletion on the scale of the entire major arc.

In the next step, we intended to incorporate a second independent variable, referred to as the Contact Zone (CZ), into our model. The CZ variable was coded for each 100×100 bp cell as 1 within the zone (6-9kb and 13-16kb) and 0 for regions outside of this zone.

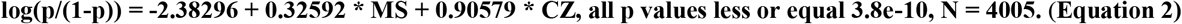

Our results indicate that the presence of a contact zone has a significant and positive impact on the probability of deletions. By using standardized variables in the equation, we can compare the coefficients and determine that the contact zone variable affects the odds of probability three times stronger (0.91 versus 0.33) than microhomology similarity.

In order to pinpoint the exact location of the macromolecular contact zone, we conducted additional data-driven analyses. Instead of using the contact zone variable, we introduced a variable with the Euclidean distance from the contact point to each cell of our contact matrix. We hypothesized that there is one contact point that, in conjunction with the microhomology score, most effectively explains the distribution of deletions (as depicted on scheme in Fig 2A). To test this hypothesis, we ran 4005 logistic regressions, each with a different contact point as the center of all 4005 cells in our matrix (all cells excluding the diagonal zone). We observed that the strongest contact point (i.e. contact point corresponding to the model with the minimal Akaike information criterion, AIC) has the coordinates of 7550 bp as 5’ and 15150 bp as 3’. Plotting the heatmap with AIC for each contact point, we demonstrated that the data-driven contact zone was similar to our visually-derived 6-9kb vs 13-16kb contact zone (fig 2C upper right triangle). Altogether, we propose that the distribution of human mtDNA deletions is determined by both the macromolecular contact zone of the single-stranded major arc and the local microhomologies between DNA regions (Fig 2C).

### 3 Single-stranded major arc of mtDNA can be folded into a large-scale loop due to DNA properties such as inverted repeats

Single-stranded DNA can maintain its structure through various factors, such as specific proteins like SSB. However, when single-stranded DNA is not covered by any proteins, it may become more susceptible to structural changes and deletions. Given that deletions in mtDNA occur infrequently (and might be associated with fluctuations in abundance of SSB) and most likely during the dynamic process of mtDNA replication when single-stranded DNA is not fully covered by protective proteins, we sought to investigate the spatial structure of the single-stranded major arc of mtDNA: to study the shape that the major arc would take based solely on its DNA properties.

To reconstruct the spatial structure of the single-stranded parental heavy chain of the major arc of mtDNA, we performed an *in silico* folding using Mfold (see Methods). Using the Mfold output obtained for the single-stranded DNA molecule of the parental heavy chain, we derived a contact matrix as the number of hydrogen bonds between two DNA regions (Fig 2D, bottom-left triangle). We observed an interesting pattern in the contact matrix: the pattern, which is a diagonal from the lower left to the upper right part of the matrix, overlapped with the contact zone between 6-9 kb and 13-16 kb. This cross-like contact matrix graph resembles bacterial Hi-C data [35], and suggests that the single-stranded heavy chain forms a hairpin-like structure, with a center close to 11 kb and large-scale stem formed by regions that are aligned with each other, such as 9.5 kb in front of 11.5 kb, 8.5 kb in front of 12.5 kb, 7.5 kb in front of 13.5 kb, and the strongest contact found at 6.5 kb in front of 14.5 kb (bottom-left triangle of Fig 2D).

The global secondary structure of the single-stranded DNA is thought to be maintained by microhomologies including inverted repeats, which can hybridize with each other to form stems. The Mfold program uses the abundance and similarity of different inverted repeats to reconstruct the shape of the single-stranded DNA. To validate the results obtained using Mfold, we correlated the density of the inverted repeats in 100 bp windows with the corresponding contacting densities of the in silico Mfold folding matrix (bottom-left triangle of Fig 2D). We observed a positive correlation between the two variables (Spearman's rho = 0.05, p = 0.0017, N= 4005, diagonal elements were removed from the analyses), which confirms that both Mfold predictions and the density of inverted repeats show a similar trend.

The in-silico folding of a very long (~10kb) single-stranded DNA molecule, as used to generate the result in Fig 2D (bottom-left triangle), may have computational limitations and artificially force the origin of the hairpin-like structures. To avoid this potential problem, we split the major arc into short (100 bp) windows, folded all pairwise combinations, estimated Gibbs Energies for each pair, and finally reconstructed the fine-scale contact matrix (upper-right triangle in Fig 2D). The fine-scale contact matrix graph shows several stripes corresponding to the strongest contacts (three horizontal lines with ordinate equals 6100, 6900 and 7900 and one vertical with abscissa equals 15000), and the intersection of these lines overlaps well with a contact zone. Altogether, the in silico folding approach supports the existence of a contact zone between 6-9 kb and 13-16 kb of mtDNA based on pure properties of single-stranded DNA.

### 4 The contact zone describes dynamics of deletions occurred during healthy aging

A recent study used an ultrasensitive method to uncover around 470,000 unique deletions in the human mtDNA [36]. Bioinformatic analysis of this dataset revealed three principal components, describing the main properties of deletions: (i) disease-versus healthy-associated deletions, (ii) located in the minor or major arcs; (iii) young or old age at the time of biopsy. Deletions with high scores on the third principal component were found by the authors [36] to be (a) associated with advanced age, (b) located primarily within the major arc of the mtDNA, (c) having high microhomology similarity between breakpoints and (d) were located in a specific manner in the major arc, where breakpoints near origins of replication and in the middle of the major arc were mostly penalized (figure 4C in the original paper [36]). This specific manner of deletion locations strongly resembles the contact zone derived in our study. This similarity assumes that formation of age-related deletions - the most common deletions in the human population - is driven mainly by the contact zone. Indeed, using the principal component analysis metadata provided by the authors, we confirmed that the scores of the third principal component of the major arc were significantly higher for bins located within the macromolecular contact zone as compared to bins located outside the contact zone (p-value < 4.48^−13^, Mann-Whitney U test, Supplementary Fig 2). This shows that the spatial structure of single-stranded mtDNA, and particularly the contact zone, plays an important role in the formation of healthy age-related deletions. It is important to emphasize also, that the mechanism of origin of this class of deletions, was proposed to occur as a primer slippage during the asynchronous strand displacement mtDNA replication [36], thus completely corroborating our findings that spatial structure of the single-stranded parental heavy chain is of great importance for the deletion formations (Fig 1, Fig 2, see also an integral scheme of the deletion formation in Fig 3).

**Figure 3.**
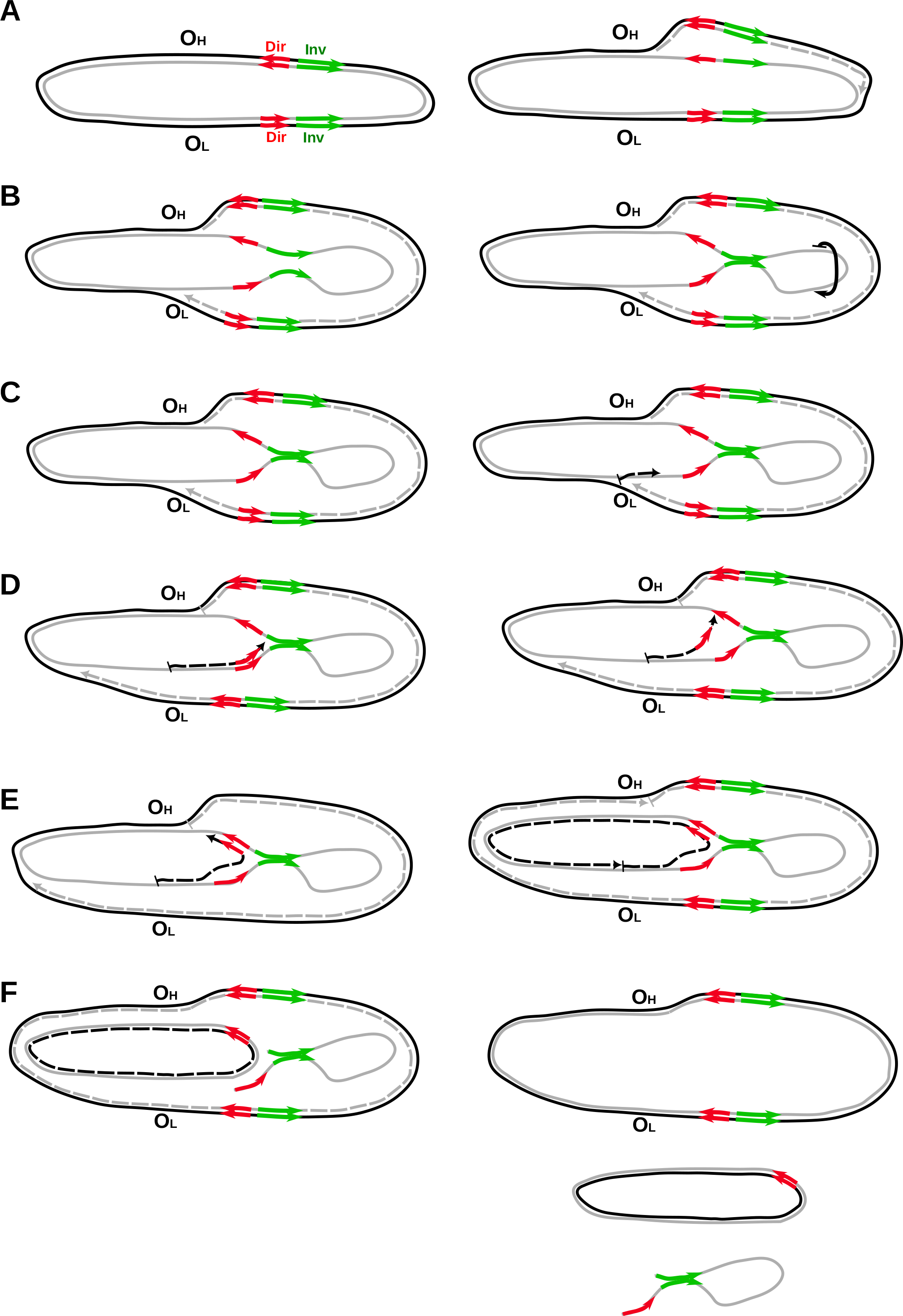
Integral scheme of the origin of mtDNA deletions. The parental heavy strand is marked by a grey line, while the parental light strand is marked by a black line. Daughter strands are indicated by dotted lines. Direct nucleotide repeats are marked by red arrows, and inverted nucleotide repeats are marked by green arrows. The origin of replication of the heavy strand (O_H_) and the origin of replication of the light strand (O_L_) are labeled. (A) At the start of replication, the daughter heavy strand (dotted gray line) is replicated on the template of the parental light strand (black line), and the replication fork begins to move from O_H_ towards O_L_. (B) During this time, the parental heavy strand remains single-stranded (ssDNA) for a significant amount of time, and different types of microhomology, including inverted repeats (green arrows), can fold the spatial structure. If ssDNA forms a stem, it may rotate (as indicated by the black arrow), potentially bringing direct repeats even closer together. (C) Once the first replication fork reaches OL, the second one begins replicating the daughter's light strand (dotted back line) on the template of the parental heavy strand. (D) The second replication fork stalls near the stem initiated by the inverted repeats. If the stalling time is sufficiently long and the spatial structure of ssDNA cannot be resolved, the new-synthesized daughter chain dissociates partially and re-aligns with the second arm of the direct repeat. (E) This allows the replication fork to continue replication. (F) Replication of the daughter light strand is finished, and the ssDNA loop region disappears either in the second round of mtDNA replication (See figure 5f in [29]) or when mtDNA with deletions is repaired, and ssDNA is degraded.

### 5 The double-sranded major arc of mtDNA may be also folded into large-scale loop: Hi-C data support the “infinity symbol” model

According to a recent report [29] and our results (Fig 1, Fig 2, Fig 3) mtDNA deletions are believed to originate during mtDNA replication when a long stretch of the parental heavy strand remains single-stranded. However, double-stranded mtDNA can also adopt a shape similar to that of single-stranded mtDNA, where both origins of replication - heavy and light - are located proximally in the contact zone. This proximity may facilitate the regulation of replication through direct crosstalk between the two origins of replication. To test the spatial structure of double-stranded mtDNA, we used the publicly available high-density contact matrix from Hi-C experiments on human lymphoblastoid cells with an available resolution of at least 1 kb [37]. We observed high-density contacts between 0-1 kb and 15-16.5 kb, which likely reflects the circular nature of mtDNA (nucleotides with positions 1 and 16569 are neighbour nucleotides). This confirms that the spatial reconstruction of mtDNA from the Hi-C data is reliable and meaningful (Supplementary figure 3, ovals mark the contacts reflecting the circular nature of mtDNA). Next, we observed the second contact which was within the major arc and strongly reminded the contact zone: 6-9 kb versus 13-16 kb (Supplementary figure 3, dotted squares mark the potential contact zone). This mtDNA contact zone suggests that the double-stranded major arc can adopt also a loop-like shape, and the entire double-stranded mtDNA may resemble an "infinity symbol" with contact zones between positions 6-9 kb and 13-16 kb.

To check the robustness of the publicly available mtDNA Hi-C matrix (31), we additionally obtained six Hi-C contact matrixes of mtDNA derived from olfactory receptors of controls and Covid patients [38]. The analysis of these contact matrices, despite low coverage and technical noise, supported the existence of contacts between positions 0-1 kb and 15-16.5 kb, reflecting the circular nature of mtDNA, as well as contacts between positions 6-9 kb and 13-16 kb, supporting the "infinity symbol" model (as shown in Supplementary Figure 4). No significant differences were observed between COVID-19 patients and controls (Supplementary Figure 4). However, further high-resolution Hi-C studies focused on mtDNA are needed to further clarify the shape of double-stranded mtDNA

## Discussion

We proposed that the human single-stranded heavy chain of mitochondrial major arc has a hairpin structure with a contact zone between 6-9 kb and 13-16 kb, which affects the mtDNA deletion spectrum (Fig 1, Fig 2, Fig 3). Our findings indicate that the formation of deletions is influenced not only by the DNA similarity between the breakpoint regions but also by the spatial structure. These results support the replication slippage mechanism where the nested pattern of direct and inverted repeats (hereafter DIID: Direct Inverted Inverted Direct) can lead to the formation of deletions [29] [28].

Our hypothesis is supported by several experiments. The first notable experiment was conducted on the mtDNA of Nematomorpha, which has high levels of perfect inverted repeats of significant length [39]. The study found that inverted repeats can form hairpins and influence DNA replication in PCR amplification. The study showed that the DIID pattern disappeared during PCR, suggesting that shorter products (mtDNA with deletion) are likely a result of PCR jumping facilitated by the presence of direct repeats flanking the hairpin. This demonstrates that the DIID pattern is indeed highly mutagenic and can lead to deletion formation (See supplementary Figure 2 in the paper [39]). A second experiment on human mtDNA showed that replicative polymerases can cause deletions through copy-choice recombination between direct repeats and that this effect is enhanced by secondary structures [29], which are maintained by inverted repeats. Third, a previous study has shown that short-sequence homologies (i.e. direct repeats) play a role in deletion formation in bacteria [28]. Furthermore, the hotspot of deletions was found to be characterized by a secondary structure, maintained by inverted repeats [28], which closely resembles the fragile DIID pattern proposed in our study. Fourth, the stalling of the DNA polymerase in the vicinity of the common repeat of human mtDNA has been demonstrated as a prerequisite of the common deletion formation [40]. According to our proposed mechanism, this stalling can be initiated by the contact zone.

Future experiments may shed light on the topological insight of the mtDNA deletion formation. First, until now, there has been no experimental reconstruction of the spatial structure of a single-stranded parental heavy chain of the major arc during human mtDNA replication. This would be a direct and important experiment to be performed. Second, our model suggests the maintenance of a contact zone, even in a double-stranded mtDNA, can put in close spatial proximity two origins of mtDNA replication (heavy and light) that will facilitate their cross-talk and co-regulation. This can be achieved by high-quality mitochondrial HiC. Third, since our model helps to rank the mutagenic potential of direct repeats according to their location (3-times higher mutability of direct repeats in the contact zone), it is testable in future experiments with other human haplogroups and other species. Fourth, our topological model can be extended to a minor arc, where deletions, although rare enough, still happen.

A deeper understanding of the deletion formation process opens a possibility of predicting mtDNA deletion spectra (distribution of different types of deletions) and deletion burden (total fraction of mutated versus wild-type mtDNA) based on mtDNA sequence and thus aids in the uncovering of haplogroup-specific mtDNA disease risks. For example, in the haplogroups with the disrupted common repeat (D4a, D5a, N1b1), we expect to observe the decreased somatic mtDNA deletion burden [23] and, consequently: postponed aging [41,42], decreased rate of neurodegeneration [6], frequency of Parkinson diseases [5], skeletal muscle myopathies [7,8], extraocular muscle weakness [9], rate of mitochondrial aging in HIV patients [43] and rate of early-onset “mtDNA deletion syndromes” classically consisting of Kearns-Sayre syndrome (KSS), Pearson syndrome and progressive external ophthalmoplegia (PEO) [13,14]. Haplogroup-specific mtDNA secondary structures, which can be obtained experimentally or computationally, can add an additional factor explaining the mtDNA deletion risks and associated variation in mtDNA-related phenotypes. Because of the high mutagenicity of spatially proximate mtDNA regions, we expect that mtDNA secondary structure may play an important role in the explanation of haplogroup-specific risks of encephalomyopathies and other human phenotypes [44].

The possibility of predicting mtDNA deletion burden and spectrum based on mtDNA sequence would offer an important step forward for mitochondrial medicine. Haplogroups with low expected deletion burden can provide a preferable donor mtDNA in mitochondrial donation [45–47] and mitochondrial transplantation [48]; [49] approaches. Additionally, a predicted haplogroup-specific spectrum of deletions can potentially help to establish a way of using of targeted systems for the elimination of expected deletions in neurons and muscle cells of aged individuals ([50,51]; [52,53]; [54]).

It would be beneficial to use comparative species data to extend our hypothesis to the evolutionary scale and demonstrate that DIID patterns increase the occurrence of deletions in the mtDNA of all species. Initially, it was reported that the mammalian lifespan negatively correlates with an abundance of direct repeats in mtDNA [25,26], suggesting that direct repeats lead to the formation of mtDNA deletions, limiting the lifespan. Later, it was found that inverted repeats have an even stronger negative correlation with mammalian lifespan [55]. Recently, a strong positive correlation was observed between the abundance of direct and inverted repeats [34]. Overall, our study suggests that the abundance of both direct and inverted repeats affects the amount of fragile DIID patterns, which are expected to be the best predictors of the somatic deletion burden and mammalian lifespan. Annotation of DIID patterns in mtDNA of all mammals or all vertebrates would open up an exciting potential direction for future research in the field.

## Methods

### Distribution of the centers

For each deletion from MitoBreak in the major arc (5781-16569), its midpoint was found. Next, each of the real deletions was moved randomly within the major arc, and their midpoints were also obtained. For the observed means of the observed deletions and randomly simulated ones, the corresponding standard deviations were obtained and compared.

### Hi-C mtDNA contact matrix

The publicly available mtDNA matrix was visualized using Juicebox. http://aidenlab.org/juicebox/?juiceboxURL=http://bit.ly/2Rmz4wy. The corresponding paper describing the methodology of obtaining Hi-C data derived from the human lymphoblastoid cell line is by Rao et al [37]. Additionally, we obtained six Hi-C mtDNA contact matrixes from olfactory receptors of covid patients and controls. Details of the in situ Hi-C protocol, as well as bioinformatics analyses, are described in the original paper [38]. Matrices were visualized using Juicebox [56].

### In silico folding

We used the heavy chain of the reference human mtDNA sequence (NC_012920.1) since it spends the most time being single-stranded according to the asymmetric model of mtDNA replication [29]. Using Mfold [57] with parameters set for DNA folding and a circular sequence, we constrained everything but the major arc from forming base pairs. We obtained the global (genome-wide) secondary structure, which we then translated into the number of hydrogen bonds connecting our regions of interest (100 bp windows for the analyses and visualization).

Next, within the single-stranded heavy chain of the major arc, we defined 100 bp windows and hybridized all potential pairs of such windows using ViennaRna Package 2 [58]. Obtained Gibbs Energies for each pair of such windows was used as a metric of a strength of a potential interaction between two single-stranded DNA regions.

### The density of inverted/direct repeats

For each pair of 100 bp window, we estimated the number of nucleotides involved in at least one inverted/direct degraded repeat. The corresponding repeat should have one arm located in the first window and another arm located in the second window. All degraded (with the maximal level of imperfection of 80%) repeats in the human mtDNA were called using our algorithm described previously [34].

### Clusterization of deletions

For clusterization, we used all MitoBreak [31] deletions from the major arc. We used 5’ and 3’coordinates as input for a hierarchical density-based clustering algorithm (python hdbscan v0.8.24). DBSCAN is a well-known algorithm for probability density-based clusterization, which detects clusters as regions with more densely located sample data as well as outlier samples. The advantage of this method is soft clustering. We variated cluster density parameters in order to ensure cluster stability and found that cluster formations stay relatively stable for a wide range of parameters. Thus, DBSCAN produces a robust set of clusters, producing additional evidence for regions with elevated deletion rates. We also performed affinity propagation clustering [59] as a data exploration experiment, which also yields robust clustering.

<u>*Perfect direct repeats of the human mtDNA:*</u> the list of the perfect direct repeats with a length of 10 or more base pairs was used from our algorithm described in Guo et al [23].

### Realized and non-realized direct degraded repeats

We used our database of degraded mtDNA repeats [34] with a length of 10 bp or more and a similarity of 80% or more. We took into account only direct repeats with both arms located in the major arc. We grouped repeats with similar motifs into clusters so that each considered cluster should contain at least three arms of the repeat, and at least one deletion should be associated with two of them. We additionally restricted our subset of clusters, considering only non-realized repeats as pairs of arms, where at least one of them (the first or the second) is the same as in realized repeat. Visually in Fig 2D, it means that within each cluster, we compare realized repeats (red dot) with non-realized ones (grey dot) located on the same horizontal (the same Y coordinate) or vertical (the same X coordinate) axis. We got 618 clusters like this.

### Pairwise alignments for microhomology matrix

A measure for the degree of similarity between segments of the major arc was obtained by aligning small windows of the mitochondrial major arc sequence with each other. We sliced the mitochondrial major arc sequence into 100 nucleotide pieces and aligned them against each other using EMBOSS Needle [60] with default parameters (match +5, gap open - 10, gap extend - 0.5), parsed out the alignment scores, thus obtaining data for the matrix of microhomology.

## Code and data availability

All data sets and scripts used in the manuscript are available on GitHub https://github.com/Aragret/ComparativeGenomics, with the key scripts, deposited within the branch HumanMtDnaDeletions, being MitoBreakDeletionsAndOrlovRepeats.R, MitoBreakDeletionsDistribution.R, RealizedVsNonrealizedDeletions.R, SlipAndJump.R and MitoBreakDeletionsAndInteractionOfDirectAndInvertedVictorRepeats.R.

## Author contributions

KP designed the study, KG, A.A.M, A.G.M, V.SH, and K.P. performed the main statistical analyses, K.U derived in silico folding and microhomologies, all authors contributed towards the writing of the manuscript.

## Acknowledgments

Desing of the study by K.P. was supported by Russian Science Foundation [No. 21-75-20143]. Main statistical analysis by K.G. and V.S. was supported by the Russian Science Foundation [No.21-75-20145]. Data management by I.M. was supported by the Russian Science Foundation [No. 21-75-10081]. E.O.T. was supported by a Ph.D. fellowship from the Austrian Science Fund FWF (DOC 33-B27). We acknowledge Filipe Pereira and Joana Damas for discussion of the MitoBreak database, Maria Falkenberg for the discussion of the potential structure of mtDNA and Nariman Battulin for the discussion of mtDNA Hi-C data. We acknowledge Scott Lujan and Bill Copeland for providing the metadata of the principal component analysis from their paper, Maxim Ri for editing and improving the manuscript. Contributions from JBO supported by grant K23DC019678 from the National Institute on Deafness and Other Communication Disorders and the National Institutes of Health as well as through grant UL1TR001873 from the National Center for Advancing Translational Sciences, National Institutes of Health. The content is solely the responsibility of the authors and does not necessarily represent the official views of the NIH. Also this work was supported by Priority 2030 at the Immanuel Kant Baltic Federal University.

**Supplementary Figures:**

Supplementary Figure_1: Distribution of the perfect direct repeats (red dots) and deletions from MitoBreak (grey dots) in the major arc.

Supplementary Figure 2: The third principal component scores, associated with aging-related deletions of healthy samples from a paper [36]. The contact, marked by the pink square, is characterized by the increased scores (p-value < 4.48^−13^, Mann-Whitney U test).

Supplementary Figure 3: Hi-C contact matrix of mtDNA obtained from the human lymphoblastoid cells. Dotted squares mark the potential contact zones. Ovals mark the contacts, emphasizing the circularity of mtDNA.

Supplementary Figure 4: Hi-C contact matrix of mtDNA obtained from the human olfactory epithelium autopsies. The top row represents two contact matrixes from covid patients, middle and bottom rows represent the contact matrices from controls. Solid white squares mark the potential contact zone. Dotted white rectangles mark the contacts, emphasizing the circularity of mtDNA.

## Notes

http://aidenlab.org/juicebox/?juiceboxURL=http://bit.ly/2Rmz4wy

https://github.com/Aragret/ComparativeGenomics/

https://github.com/Aragret/ComparativeGenomics/blob/HumanMtDnaDeletions/Body/4Figures/Figure2final_23072020WithOvalsKP1.pdf

https://github.com/Aragret/ComparativeGenomics/blob/HumanMtDnaDeletions/Body/4Figures/Fig2_transparency_20230126-KG.pdf

https://github.com/Aragret/ComparativeGenomics/blob/HumanMtDnaDeletions/Body/4Figures/Fig3_20230120-KP.pdf

https://github.com/Aragret/ComparativeGenomics/blob/HumanMtDnaDeletions/Body/4Figures/SupplFig2_19092020RedDotsVSh.pdf

https://github.com/Aragret/ComparativeGenomics/blob/HumanMtDnaDeletions/Body/4Figures/CopelandThirdComponentPca2.R.svg.pdf

https://github.com/Aragret/ComparativeGenomics/blob/HumanMtDnaDeletions/Body/4Figures/SupplFig1HiC_31052022.pdf

https://github.com/Aragret/ComparativeGenomics/blob/HumanMtDnaDeletions/Body/4Figures/SupplFig2HiC_20230118-KG.pdf

### Competing Interest Statement

The authors have declared no competing interest.

### Summary of Updates

We paid special attention to the logic and presentation of the material and completely changed the order of the sections in the results (we started from the data-driven contact zone obtained on ssDNA and extended it at the end to Hi-C). For example, our new section 1 corresponds to old sections 3 and 4; new section 2 corresponds to old 5; 3 - 2; 4 - 6, and 5 - 1. Also, we significantly improved figure 2, and all supplementary figures, and we added figure 3 with an integral scheme of the proposed mechanism of deletion formation. Altogether, we believe, our paper is much more clear and more straightforward now. Instead of including additional analyses on HiC data, the results related to HiC have been toned down and are now presented as a suggestive, rather than conclusive, section. The main conclusion of the paper, which focuses on the folding of the single-stranded heavy chain, remains valid regardless of the results from the HiC data. The HiC data is included primarily to demonstrate the need for further experimental research in this area and the possibility that even double-stranded mtDNA may have a secondary structure similar to the infinity symbol. We realized that the oval shape of the contact zone is interesting and potentially biologically meaningful, as seen in other figures, such as Figures 2A and 2B. We have discussed this phenomenon further in the third paragraph of the first results section. First of all, we would like to emphasize that currently, this analysis moved closer to the beginning of the results, and thus, it helps to establish the contact zone per se (data-driven search of the region of the contact zone). Second, exact probabilities are difficult to derive since the origin of deletions is an extremely rare process: it most likely depends on the genome of a patient, age, etc. So any estimations of the absolute probabilities of deletions would base on extremely simplified assumptions and unlikely can be biologically meaningful. This question deserves a separate follow-up, where more parameters can be included. However, to run some pilot calculations, we transformed the logit function into probabilities based on the definition of logistic regression. We tested the logarithmic regression with the coefficients from equation II given in the article.

