## Supplementary figures and images for "Risk of mitochondrial deletions is affected by the global secondary structure of the human mitochondrial genome"

### Supplementary Figure 1

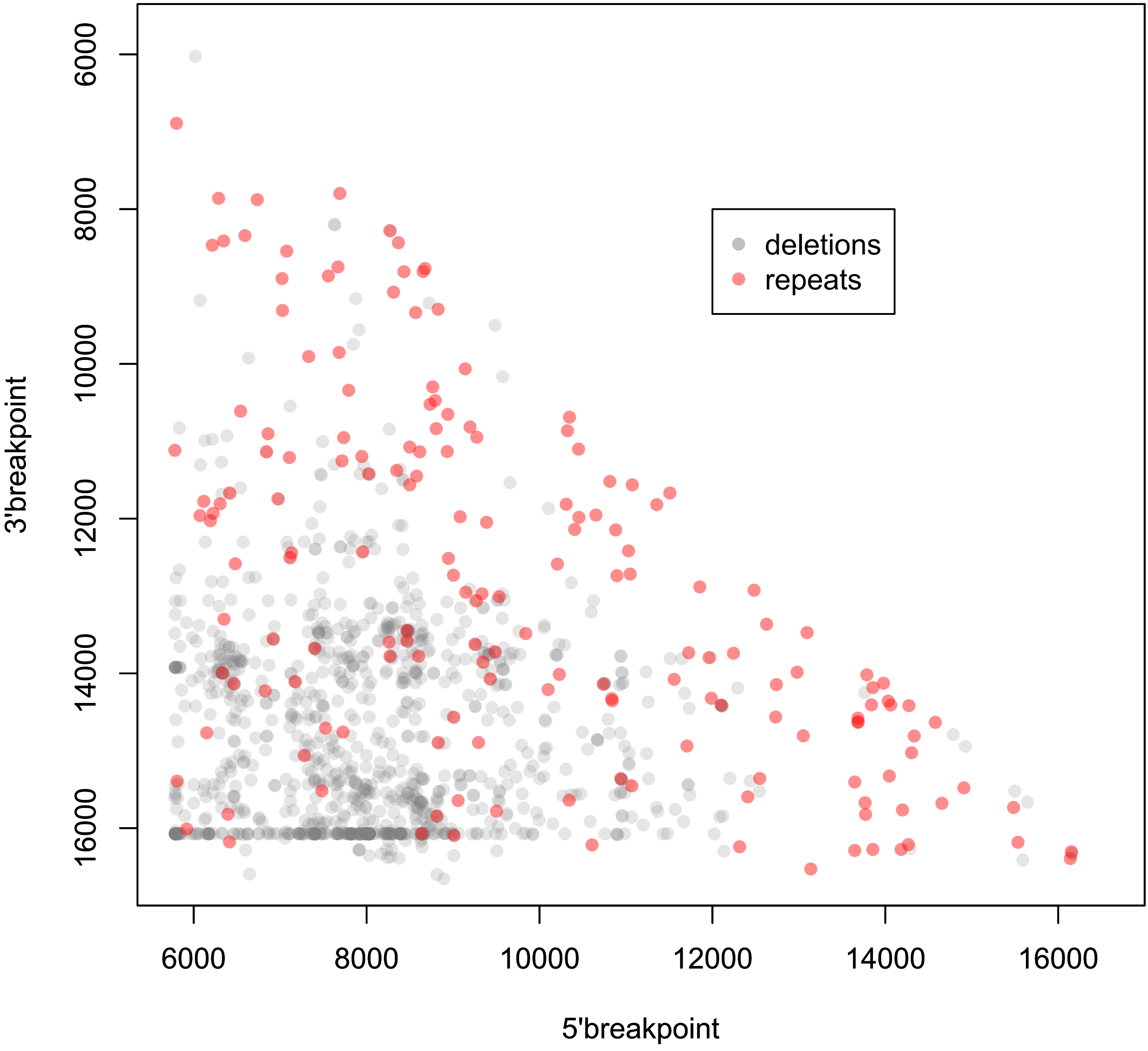

### Supplementary Figure 2

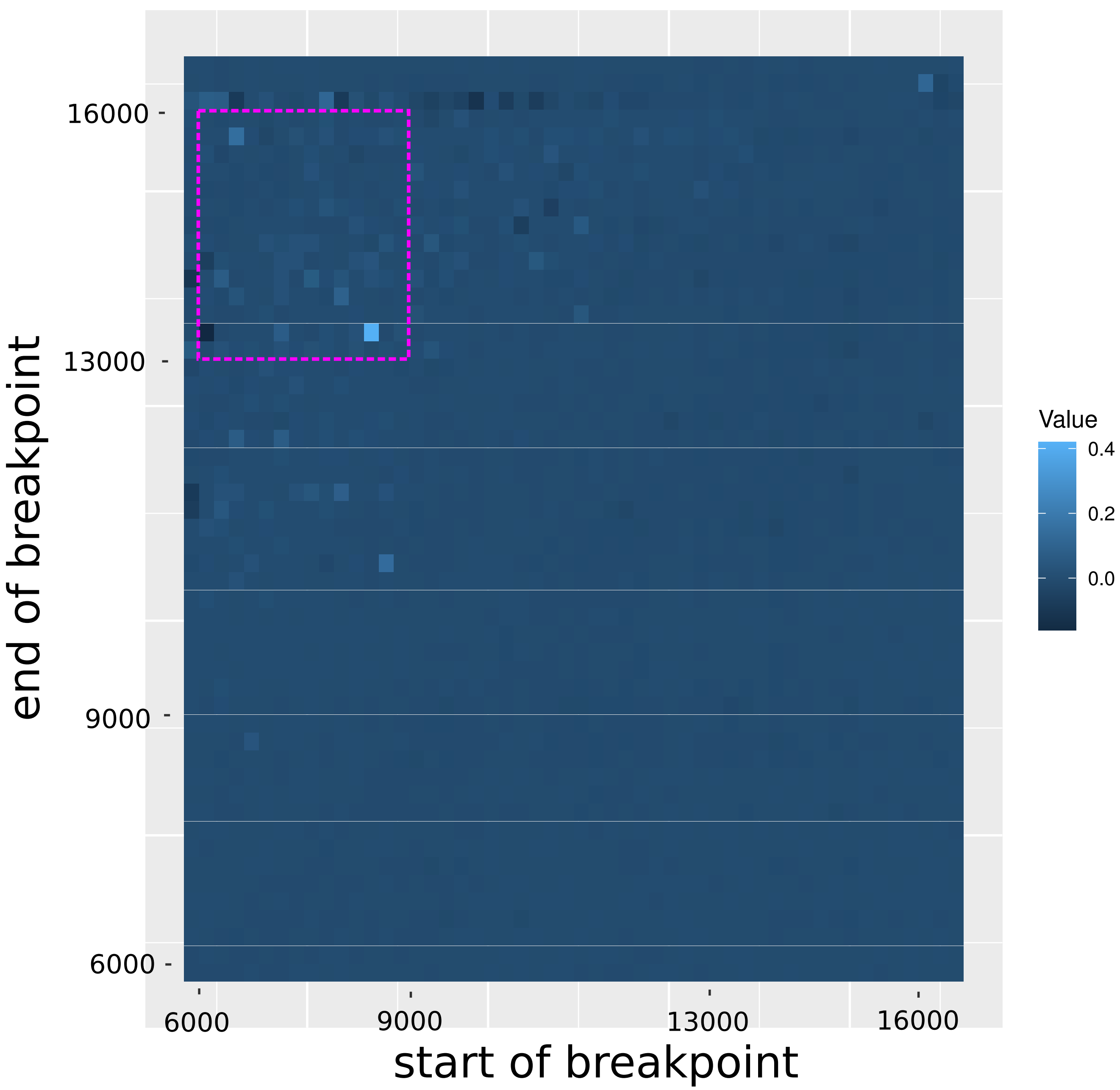

### Supplementary Figure 3

#### mtDNA Hi-C contact matrix

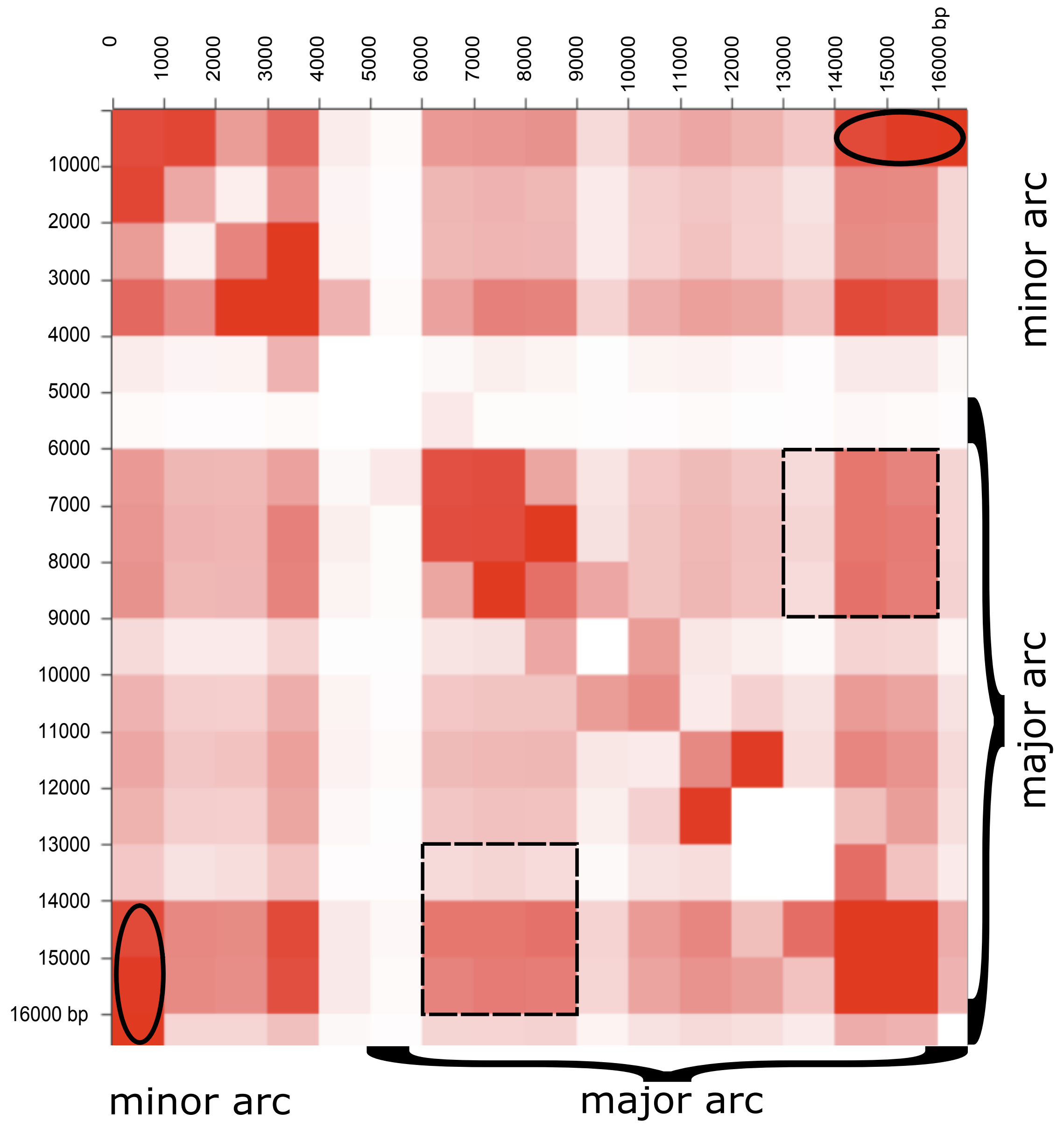

### Supplementary Figure 4

#### mtDNA Hi-C contact matrix

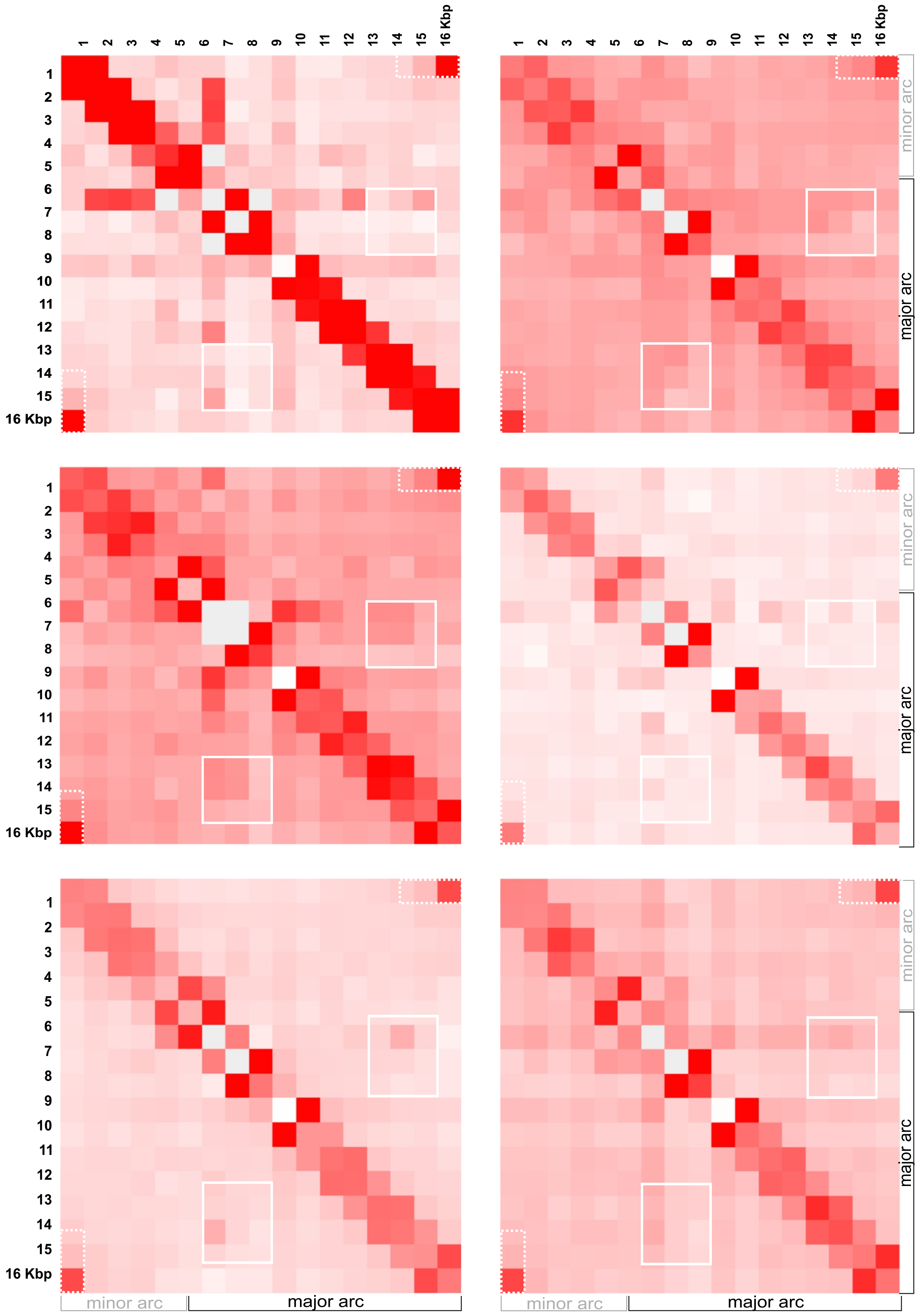
